# Graph Neural Networks for Z-DNA prediction in Genomes

**DOI:** 10.1101/2022.08.23.504929

**Authors:** Artem Voytetskiy, Alan Herbert, Maria Poptsova

**Affiliations:** *Bioinformatics Laboratory, HSE University*, Moscow, Russia; *InsideOutBio*, Charlestown, MA, USA

**Keywords:** Z-DNA, flipons, non-B DNA, Recurrent Neural Network, Graph Neural Network, Graph Convolutional Network Graph Attention Network, GraphSAGE

## Abstract

Deep learning methods have been successfully applied to the tasks of predicting functional genomic elements such as histone marks, transcriptions factor binding sites, non-B DNA structures, and regulatory variants. Initially convolutional neural networks (CNN) and recurrent neural networks (RNN) or hybrid CNN-RNN models appeared to be the methods of choice for genomic studies. With the advance of machine learning algorithms other deep learning architectures started to outperform CNN and RNN in various applications. Thus, graph neural network (GNN) applications improved the prediction of drug effects, disease associations, protein-protein interactions, protein structures and their functions. The performance of GNN is yet to be fully explored in genomics. Earlier we developed DeepZ approach in which deep learning model is trained on information both from sequence and omics data. Initially this approach was implemented with CNN and RNN but is not limited to these classes of neural networks. In this study we implemented the DeepZ approach by substituting RNN with GNN. We tested three different GNN architectures – Graph Convolutional Network (GCN), Graph Attention Network (GAT) and inductive representation learning network GraphSAGE. The GNN models outperformed current state-of the art RNN model from initial DeepZ realization. Graph SAGE showed the best performance for the small training set of human Z-DNA ChIP-seq data while Graph Convolutional Network was superior for specific curaxin-induced mouse Z-DNA data that was recently reported. Our results show the potential of GNN applications for the task of predicting genomic functional elements based on DNA sequence and omics data.

**Availability and implementation:** The code is freely available at https://github.com/MrARVO/GraphZ.

## I. Introduction

Graph Neural Network approaches are applicable to any data that can be represented as a graph, as shown by previous applications of GNN that leverage many bioinformatic datasets available. The most successful implementations are for prediction of drug-target, ligand-protein or protein-protein interactions (see [1] for review).

Later graph representation learning has been extended to genomics. A graph neural representation of RNA-protein interactions based on an adjacency matrix containing basepairing probabilities was used to predict RNA structure [2]. A Graph Convolutional Network used the k-mer co-occurrence and relationships binding to learn the underlying k-mer graph to successfully model DNA-protein interactions [3]. A Graph Attention Network (GAN) was applied to Hi-C data and gene co-expression networks to produce a graph representation of the gene contact network to predict chromatin organization in cells and reveal importance of gene contacts with other genes in the neighborhood [4].

The application of GNN to single cell RNA-seq analysis has been highly successful in learning the cell–cell relationships that drive biological responses. In this approach, the gene expression matrix from scRNA-Seq experiments is first processed with graph autoencoder network such as that proposed in [5]. The topological graph embeddings are then recovered to reconstruct the cell graph [6].

Graph neural network for protein function prediction was implemented in DeepGraphGO that used combination of protein sequence and high-order protein network data [7]. The GNN approach combining sequence 3D-genome organization with other information also allowed better prediction of epigenetic state than current methods based on graph convolutional networks which learn node representations from local sequence and long-range interactions [8]. The incorporation of patient omics data into the graph representation learning model allowed prediction of disease outcome by a multi-level attention GNN [9].

Here we tested graph neural networks for the task of prediction Z-DNA regions, called Z-flipons [10], which have the potential to regulate many cellular processes [11] and that are involved in mendelian disease [12]. We tested performances of several graph neural network architectures - Graph Convolutional Network (GCN) [13, 14], Graph Attention Network (GAT) [15, 16] and GraphSAGE [17].

## II. Methods

### A. Data

We took the same data as it was used in the original paper of DeepZ [18]: ChIP-seq data from Shin et al. for human genome [19] that identified 385 regions of Z-DNA after filtering for black-listed regions was tested against a dataset of 1054 omics features to predict other Z-DNA forming regions in the genome (see full list in [18]). For mouse genome we took the same data as it was used for DeepZ in [20]: ChIP-seq data resulted from curaxin treatment of mouse embryonic fibroblasts that comprised 1957 regions and 874 omics features (see full list in [20]). Schema of DNA sequence and omics data representation is presented in Figure 1.

**Fig. 1.**
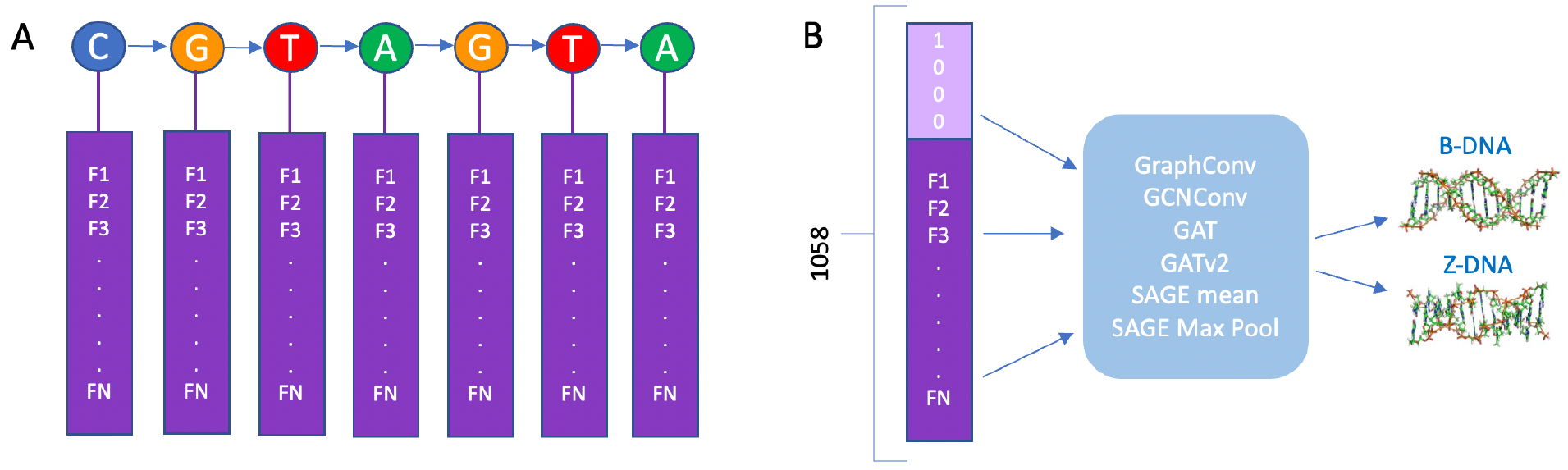
DeepZ with GNN model. (A) DNA sequence with omics features represented as vectors. (B) Schematics of DeepZ approach with different GNN architectures.

### B. Training

The whole genome was split into 5000 bp segments, and each segment was classified according to presence/absence of Z-DNA. The negative class was composed from randomly selected segments without Z-DNA. We tested several ratios of negative to positive classes. Training and test sets were split in the proportion of 4:1 with the same ratio of class balance. The training process took 25 epochs, and the hyperparameters were selected according to f1-score. With this approach we select the model with the optimal values of precision and recall, which does not suffer from overfitting.

We used 3- and 4-layes models as designated in Tables 1–4 because the performance of these models was better than of other configurations. We also trained models on data sets with different ratio of positive and negative classes. For example, model termed as 5N was trained on a data set that had 5 times more elements of negative class than the positive.

**TABLE I.** Mouse genome. Prediction on the test set.

| Model | Prec. | Recall | F1 | ROC-AUC | PR-AUC |
| --- | --- | --- | --- | --- | --- |
| **GCNConv** |  |  |  |  |  |
| 3L 5N | 0.578 | 0.374 | 0.454 | 0.868 | 0.459 |
| 3L 7N | 0.59 | 0.316 | 0.412 | 0.827 | 0.414 |
| 4L 5N | 0.558 | 0.395 | 0.462 | 0.854 | 0.447 |
| **GraphConv** |  |  |  |  |  |
| 3L 5N | **0.578** | **0.404** | **0.475** | 0.873 | 0.486 |
| 3L 7N | **0.55** | **0.419** | **0.475** | 0.865 | 0.475 |
| 4L 5N | 0.621 | 0.375 | 0.468 | 0.864 | 0.484 |
| 4L 7N | 0.588 | 0.387 | 0.467 | 0.852 | 0.468 |
| **GAT** |  |  |  |  |  |
| 3L 4N | 0.649 | 0.341 | 0.448 | 0.882 | 0.502 |
| 3L 5N | 0.63 | 0.323 | 0.427 | 0.866 | 0.469 |
| 3L 7N | 0.585 | 0.361 | 0.447 | 0.87 | 0.453 |
| 4L 5N | 0.616 | 0.357 | 0.452 | 0.872 | 0.468 |
| **GATv2** |  |  |  |  |  |
| 3L 5N | 0.598 | 0.358 | 0.448 | 0.866 | 0.468 |
| 3L 7N | 0.596 | 0.349 | 0.441 | 0.868 | 0.464 |
| 4L 5N | 0.567 | 0.388 | 0.461 | 0.856 | 0.446 |
| **SAGE** |  |  |  |  |  |
| 3L 5N | 0.588 | 0.362 | 0.448 | 0.868 | 0.46 |
| 3L 7N | 0.536 | 0.398 | 0.457 | 0.872 | 0.434 |
| 4L 7N | 0.595 | 0.353 | 0.443 | 0.868 | 0.452 |
| **SAGE max pooling** |  |  |  |  |  |
| 3L 5N | 0.566 | 0.408 | 0.474 | 0.872 | 0.476 |

**TABLE II.** Mouse genome. Whole-genome prediction.

| Model | Prec. | Recall | F1 |
| --- | --- | --- | --- |
| **RNN (DeepZ)** |  |  |  |
|  | 0.40 | 0.16 | 0.23 |
| **GCNConv** |  |  |  |
| 3L 5N | 0.224 | 0.387 | 0.284 |
| 3L 7N | 0.288 | 0.323 | 0.304 |
| 4L 5N | 0.229 | 0.421 | 0.297 |
| **GraphConv** |  |  |  |
| 3L 5N | 0.19 | 0.441 | 0.266 |
| 3L 7N | 0.118 | 0.443 | 0.264 |
| **4L 5N** | **0.253** | **0.416** | **0.315** |
| 4L 7N | 0.203 | 0.422 | 0.274 |
| **GAT** |  |  |  |
| 3L 4N | 0.232 | 0.354 | 0.28 |
| 3L 5N | 0.288 | 0.342 | 0.312 |
| 3L 7N | 0.21 | 0.37 | 0.273 |
| 4L 5N | 0.217 | 0.38 | 0.275 |
| **GAT v2** |  |  |  |
| 3L 5N | 0.238 | 0.377 | 0.292 |
| 3L 7N | 0.239 | 0.362 | 0.288 |
| 4L 5N | 0.234 | 0.416 | 0.299 |
| **SAGE mean** |  |  |  |
| 3L 5N | 0.246 | 0.382 | 0.299 |
| 3L 7N | 0.215 | 0.411 | 0.283 |
| 4L 7N | 0.232 | 0.372 | 0.286 |
| **SAGE max pooling** |  |  |  |
| 3L 5N | 0.174 | 0.432 | 0.248 |

**TABLE III.**
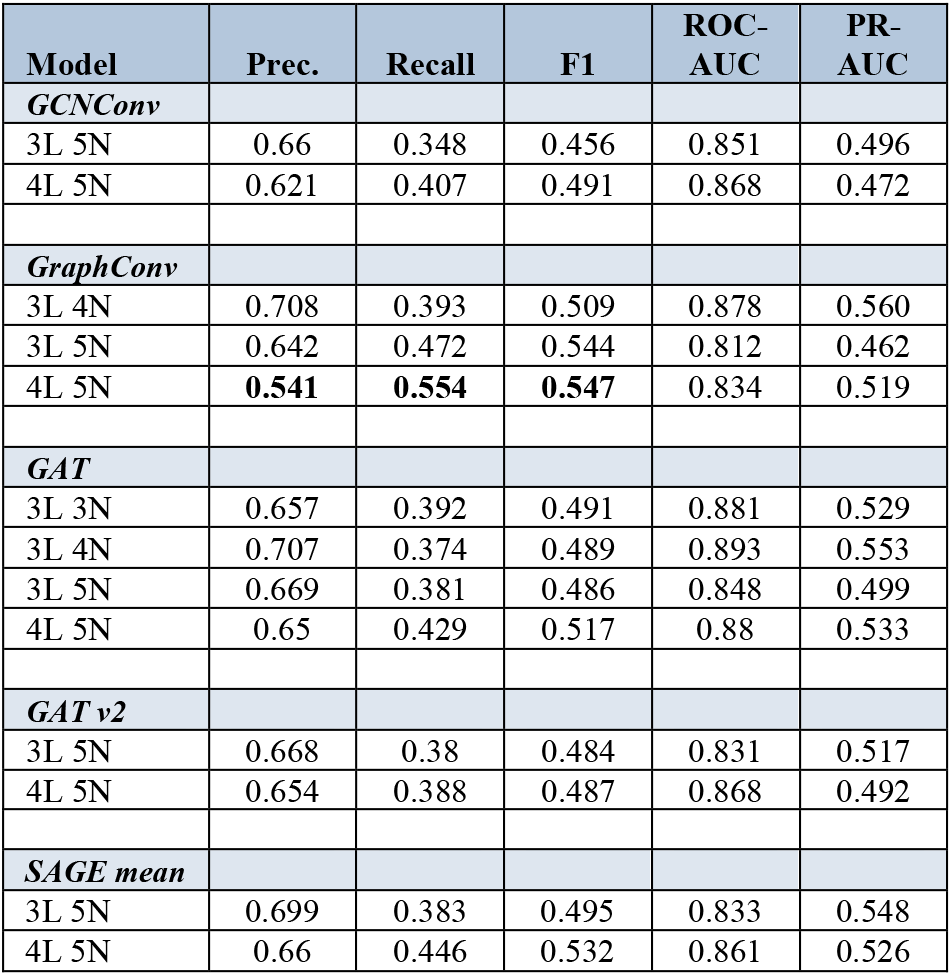
Human genome. Prediction on the test set.

| Model | Prec. | Recall | F1 | ROC-AUC | PR-AUC |
| --- | --- | --- | --- | --- | --- |
| **GCNConv** |  |  |  |  |  |
| 3L 5N | 0.66 | 0.348 | 0.456 | 0.851 | 0.496 |
| 4L 5N | 0.621 | 0.407 | 0.491 | 0.868 | 0.472 |
| **GraphConv** |  |  |  |  |  |
| 3L 4N | 0.708 | 0.393 | 0.509 | 0.878 | 0.560 |
| 3L 5N | 0.642 | 0.472 | 0.544 | 0.812 | 0.462 |
| 4L 5N | **0.541** | **0.554** | **0.547** | 0.834 | 0.519 |
| **GAT** |  |  |  |  |  |
| 3L 3N | 0.657 | 0.392 | 0.491 | 0.881 | 0.529 |
| 3L 4N | 0.707 | 0.374 | 0.489 | 0.893 | 0.553 |
| 3L 5N | 0.669 | 0.381 | 0.486 | 0.848 | 0.499 |
| 4L 5N | 0.65 | 0.429 | 0.517 | 0.88 | 0.533 |
| **GAT v2** |  |  |  |  |  |
| 3L 5N | 0.668 | 0.38 | 0.484 | 0.831 | 0.517 |
| 4L 5N | 0.654 | 0.388 | 0.487 | 0.868 | 0.492 |
| **SAGE mean** |  |  |  |  |  |
| 3L 5N | 0.699 | 0.383 | 0.495 | 0.833 | 0.548 |
| 4L 5N | 0.66 | 0.446 | 0.532 | 0.861 | 0.526 |

**TABLE IV.** Human genome. Whole-genome prediction.

| Model | Precision | Recall | F1 |
| --- | --- | --- | --- |
| **RNN (DeepZ)** |  |  |  |
|  | **0.110** | **0.270** | **0.160** |
| **GCNConv** |  |  |  |
| 3L 5N | 0.068 | 0.547 | 0.121 |
| 4L 5N | 0.037 | 0.603 | 0.07 |
| **GraphConv** |  |  |  |
| 3L 4N | 0.059 | 0.66 | 0.109 |
| 3L 5N | 0.03 | 0.757 | 0.060 |
| 4L 5N | 0.016 | 0.832 | 0.032 |
| **GAT** |  |  |  |
| 3L 3N | 0.054 | 0.6 | 0.099 |
| 3L 4N | 0.067 | 0.604 | 0.12 |
| 3L 5N | 0.062 | 0.626 | 0.112 |
| 4L 5N | 0.042 | 0.706 | 0.079 |
| **GAT v2** |  |  |  |
| 3L 5N | 0.06 | 0.593 | 0.109 |
| 4L 5N | 0.053 | 0.586 | 0.097 |
| **SAGE mean** |  |  |  |
| 3L 5N | **0.069** | **0.622** | **0.124** |
| 4L 5N | 0.043 | 0.731 | 0.082 |

### C. Graph neural networks

Schematics of DeepZ approach with different GNN architectures is presented in Figure 1B. All models were implemented with torch_geomentric that include realization of different types of layers of GNN with RMSprop optimizer and NLLLoss loss function.

Graph Convolutional Network was implemented as GCNConv [13] and GraphConv [14].

Graph Attention Network was implemented as GATConv [16] and GATv2Conv [15]. GAT architecture uses hidden layers with attention mechanisms. These layers assign to neighbors of a node individual weights that are taking into consideration during aggregation. The difference between two sub-types is that GATConv uses static attention mechanism and GATv2Conv – dynamic.

GraphSAGE was implemented as SAGEConv [17] with two different aggregation schemes: mean and max pooling. In GraphSAGE architecture instead of separate vectors for every node the training is performed for aggregation functions. Each function can gather information of different distances from the current node. This approach helps to predict nodes that represent Z-DNA forming regions not seen during the training.

### D. Benchmark model

DeepZ model was run as described in [18] for human genome data set [19] and as described in [20] for mouse genome data set [20]. Human data set comprised 1054 omics features (see full list in [18]) and mouse data set comprised 874 omics features (see full list in [20]).

## III. Results and Discussion

The results of the models’ performance on the test sets and for whole-genome prediction for human and mouse genome are presented in Tables 1–4. L signifies number of layers and N stands for the ratio between positive and negative classes. For the mouse genome data set the best model is Graph Convolutional Network that has a recall 2.6 higher than RNN (increase from 0.16 to 0.416). For the human genome, the highest F1 metric is for RNN from the initial DeepZ implementation. However, the GraphSage architecture outperforms the RNN in recall by a factor of 2.3 (0.27 vs 0.62).

Graph Attention Networks have almost the same performance as GraphSage (F1=0.120 vs F1=0.124) for the human genome and as GraphConv does for the mouse genome (F1=0.315 vs F1=0.312). In comparison, Graph Convolutional Networks can reach recall of 0.832 for human and 0.443 for mouse whole-genome predictions.

The low F-metric at the whole-genome level is explained by the fact that experimental data is incomplete due to dynamic nature of Z-DNA formation with only a subset of all possible functional Z-flipons captured from the cell type used in the experiment. However, in individual experiments, DeepZ and various GNN models predict true functional Z-DNA regions that are present in the training set, yielding considerably higher metrics for both human and mouse genomes (Tables 1–4).

The increase in performance of GNNs over RNNs and CNNs can be explained by the ability of GNN models to take into account omics features of the neighboring nodes that are encoded as node weights – making use of information that the other approaches cannot.

In our models DNA sequence is represented as a graph with each nucleotide placed at a node and omics data is assigned as features to each node (Figure 1). The advantage of graph neural network over convolution neural network is that the model learns features by aggregating features from neighboring nodes that can have many experimentally derived attributes. This approach allows extraction of relevant features from high dimensional datasets.

The three GNN architectures differ from each other by the way they learn features of neighboring nodes (Figure 2). Graph Convolutional Network is a convolutional neural network on a graph in which, instead of applying a convolution kernel, the algorithm first aggregates features from neighboring and current nodes, and then encode them depending on the result of the aggregation. As an aggregation function, GCN uses per element average and then makes a weighted sum. The distinctive property of Graph Attention Network is that feature aggregation is realized using an attention mechanism. In the original publication, the attention mechanism is implemented as an additional layer of neural network that allows aggregating data for each node using weights, thus signifying an importance of each node for its neighbors. This approach takes into account properties of neighboring nodes and improve overall results. GraphSAGE is an inductive representation learning network with an aggregation function that can use different models: average, pooling, or LSTM and an update function with concatenation. The advantage of GraphSAGE is its ability to learn from small amount of data.

**Fig. 2.**
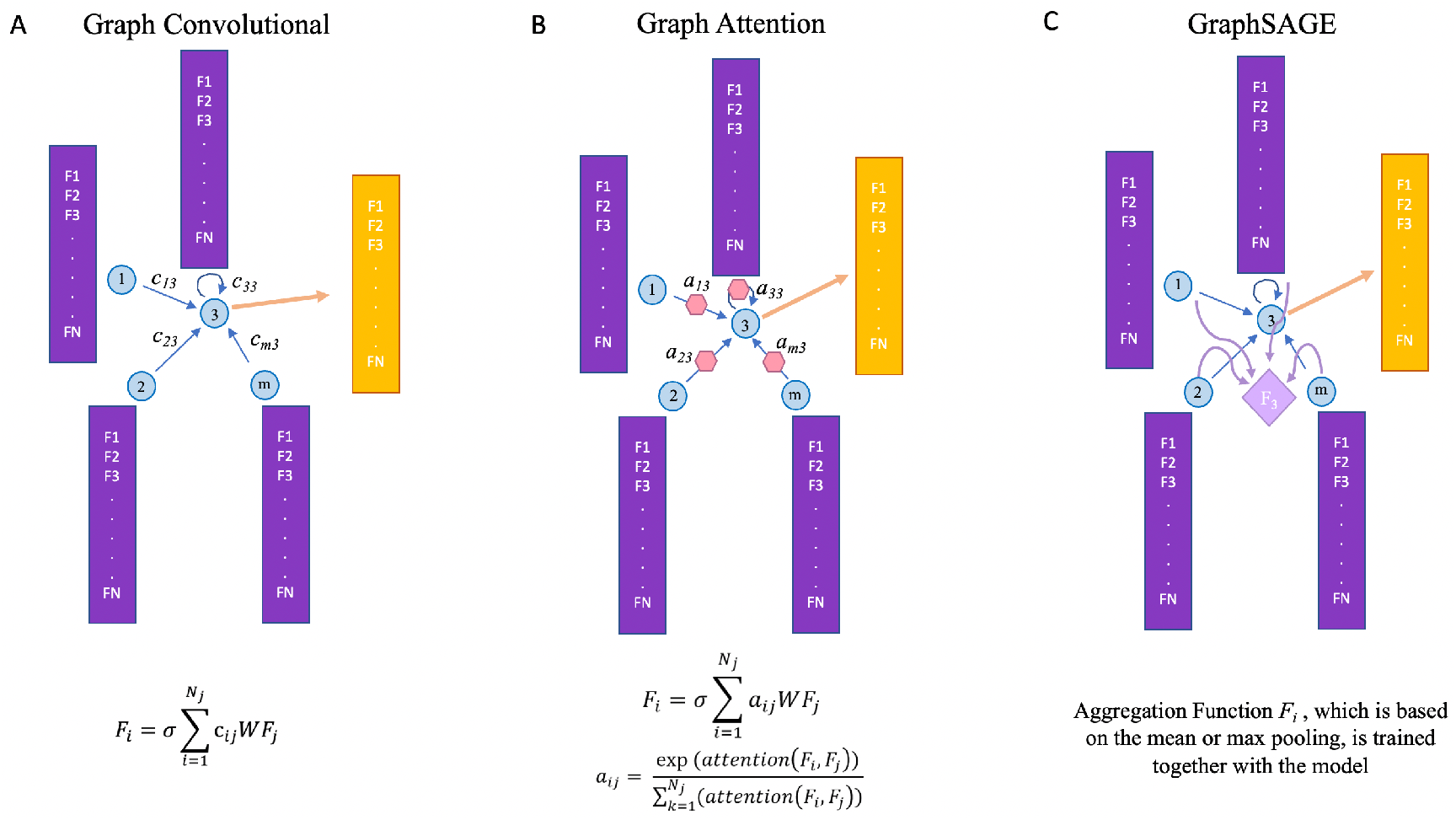
Aggregations functions for different types of GNN. A. Graph Convolutional Network B. Graph Attention Network. C. GraphSage.

## IV. Conclusion

Here we tested DeepZ approach with GNN deep learning model instead of RNN. We built and tested three major types of graph neural network modes – two types of Graph Convolutional Networks, two types of Graph Attention Networks and inductive representation learning network GraphSAGE. All three networks outperformed RNN from the initial DeepZ implementation by increasing recall in 2.3-2.6 times for whole-genome prediction. Graph Convolutional Network showed overall higher performance when compared to Graph Attention Networks and GraphSAGE, while GraphSAGE appeared better for small data sets. The presented study demonstrated potential of replacing CNN or RNN with GNN in genomic applications especially for those that rely on high dimensional omics data for training. The advantage of GNN is that it aggregate features from neighboring nodes. There is potential for improvement of GNN architecture by incorporating long-range interactions between DNA nodes into the graph representation, by using different weighing schemes that capture the correlation between features of adjacent nodes and the use of L1 metrics to reduce the model sizes.

## Acknowledgment

We thank Nazar Bekanazarov for providing DeepZ training data for mouse and human genomes. The publication was supported by the grant for research centers in the field of AI provided by the Analytical Center for the Government of the Russian Federation (ACRF) in accordance with the agreement on the provision of subsidies (identifier of the agreement 000000D730321P5Q0002) and the agreement with HSE University No. 70-2021-00139.

